# An engineered A549 cell line expressing CD13 and TMPRSS2 is permissive to clinical isolate of human coronavirus 229E

**DOI:** 10.1101/2023.07.21.545505

**Authors:** Laurensius Kevin Lie, Aleksandra Synowiec, Jedrzej Mazur, Krzysztof Pyrć

## Abstract

The lack of suitable *in vitro* culture model has hampered research on wild-type (WT) human coronaviruses. While 3D tissue or organ cultures have been instrumental for this purpose, such models are challenging, time-consuming, expensive and require extensive cell culture adaptation and directed evolution. Consequently, high-throughput applications are beyond reach in most cases. Here we developed a robust A549 cell line permissive to a human coronavirus 229E (HCoV-229E) clinical isolate by transducing CD13 and transmembrane serine protease 2 (TMPRSS2), henceforth referred to as A549^++^ cells. This modification allowed for productive infection, and a more detailed analysis showed that the virus might use the TMPRSS2-dependent pathway but can still bypass this pathway using cathepsin-mediated endocytosis. Overall, our data showed that A549^++^ cells are permissive to HCoV-229E clinical isolate, and applicable for further studies on HCoV-229E infectiology. Moreover, this line constitutes a uniform platform for studies on multiple members of the *Coronaviridae* family.

**Graphical abstract:** 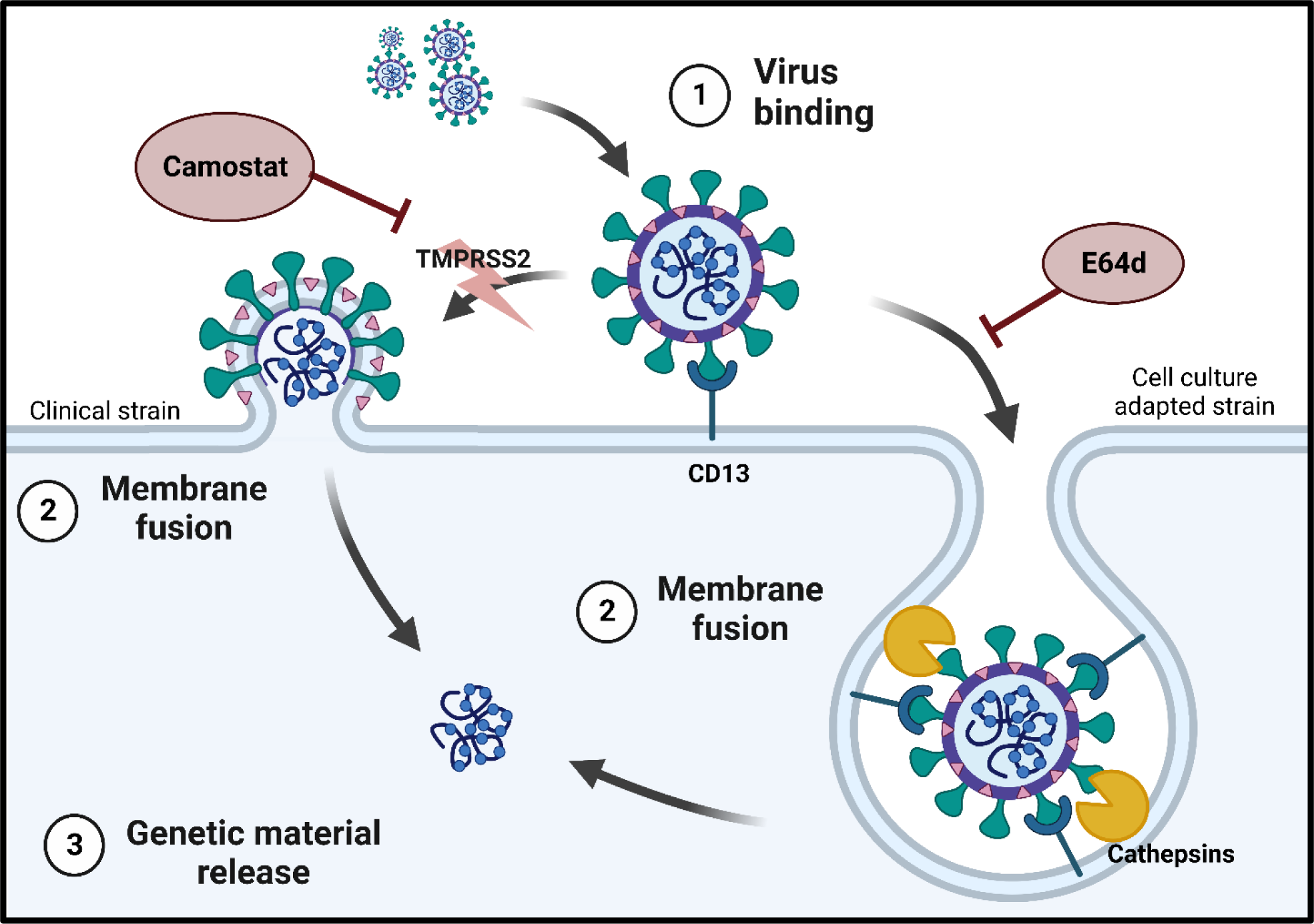

## Introduction

Human coronavirus 229E (HCoV-229E) is one of the seven coronaviruses infecting humans. First isolated in 1966, this endemic human coronavirus is a member of the genus *Alphacoronavirus* together with HCoV-NL63 and typically causes mild respiratory symptoms, though more severe symptoms have also been reported. HCoV-229E utilizes aminopeptidase N (CD13; APN) to enter permissive cells (1, 2). However, HCoV-229E can utilize at least two distinct entry pathways (3, 4). The tissue culture-adapted strain VR-740 enters the cell via the endosomal pathway and utilizes endosomal protease Cathepsin L for viral spike (S) protein activation (3). Conversely, the WT strain requires transmembrane serine protease 2 (TMPRSS2) to proteolytically prime the viral spike and bypass the endosomal entry pathway through direct membrane fusion (3, 4).

Although human stem cell-derived models of the epithelium lining the respiratory tract have been successfully used to study human coronavirus infection *ex vivo*, these models require advanced cell culture expertise, are expensive, time-consuming, and are challenging to upscale or use for high-throughput applications (5, 6). Further, WT human coronaviruses poorly propagate in cell lines and require extensive cell culture adaptation associated with accumulation of changes in, e.g., the Spike protein, which attenuates the virus *in vivo* and changes the virus phenotype (3, 7). For some species, there are no permissive *in vitro* models. HCoV-HKU1 for example, can only infect and productively replicate in the human airway epithelium (HAE) model (8).

To circumvent these challenges, there are efforts to develop novel cell models carrying the required elements and mirroring the *in vivo* infection microenvironment. For this purpose, lentiviral transduction has been popularly used to introduce transgene sequences into host cells, and integration of the transgene into the host genome ensures stable expression and inheritance of the transgene into the progeny cells (9, 10). Using lentiviral transduction, Shirato *et al.* developed HeLa cells expressing TMPRSS2 (HeLa^TMPRSS2+^) for infection with WT HCoV-229E (3), and we have also developed A549 cells expressing both angiotensin-converting enzyme 2 (ACE2) and TMPRSS2 for infection with SARS-CoV-2 (11).

Here, we report the development of a robust A549 cell line that expresses both CD13 and TMPRSS2 that supports infection by HCoV-229E clinical isolates. We investigated the replication kinetics in the abovementioned cell line, we also evaluated the production of infectious progeny and the formation of syncytia against the tissue culture-adapted strain VR-740. Finally, we mapped the entry route of these viruses in the new model.

## Materials and methods

### Cell lines

A549 (ATCC: CCL-185) and HEK 293T (ATCC: CRL-3216) cells were maintained in Dulbecco’s minimum essential medium containing 4.5 g/l glucose (DMEM, Gibco), supplemented with 5% fetal bovine serum (FBS, Gibco), 100 units/ml penicillin, and 100 µg/ml streptomycin (1% P/S, Gibco). A549^++^ cells were maintained in DMEM supplemented with 5% FBS, 1% P/S, 10 µg/ml of blasticidin S (Thermo Fisher Scientific), and 500 µg/ml of G418 (Geneticin^TM^, Gibco). MRC-5 (ATCC: CCL-171) cells were maintained in DMEM with 10% FBS and 1% P/S. A549^TMPRSS2+^ and A549^CD13+^ cells were similarly maintained in DMEM with 5% FBS, and 1% P/S, and supplemented with 10 µg/ml of blasticidin S or 500 µg/ml of G418 respectively. All cell lines were passaged every 3 to 4 days and maintained at 37°C and 5% CO_2_ unless otherwise stated. All cell lines were tested every two weeks for mycoplasma contamination either by in-house DAPI staining or LookOut^®^ Mycoplasma PCR Detection Kit (Sigma-Aldrich).

### Lentiviral Particle Production

Lentiviral vectors were produced via co-transfection of HEK293T cells using the polyethyleneimine (PEI) transfection method as described before (11). Briefly, 2 × 10^6^ HEK293T cells were seeded on a 10 cm Petri dish (TPP Techno Plastic Products AG, Trasadingen, Switzerland). Twenty-four hours later, cells were co-transfected with 500 µl of transfection mix. The mix consisted of 27 µg of PEI (Sigma-Aldrich), 9 µg of a vector carrying either the CD13/aminopeptidase N sequence (pLEX307-APN-G418 was a gift from Alejandro Chavez & Sho Iketani; Addgene plasmid #158456) or the TMPRSS2 sequence (pLEX307-TMPRSS2-blast was a gift from Alejandro Chavez & Sho Iketani; Addgene plasmid #158458). Additionally, it included 6.75 μg of the packaging plasmid psPAX2 (a gift from Didier Trono; Addgene plasmid #12260), and 2.25 μg of the envelope plasmid pMD2.G (a gift from Didier Trono; Addgene plasmid #12259), all diluted in OptiMEM (Thermo Fisher Scientific).

The medium was refreshed after 16 h and recombinant lentiviruses were harvested 24 h, 48 h, and 72 h later. The supernatant containing vector particles underwent centrifugation at 1000 × g for 5 min at 4°C, filtered through 0.22 µm PVDF membrane filters, and concentrated 20 times using Amicon Ultra-0.5 Centrifugal Filter Unit Ultracel 100 kDa cut-off (Sigma-Aldrich). Following the concentration process, we stored the resultant supernatants at a temperature of −80°C for future use.

### Generation of double-transduced (A549^++^) and single-transduced A549 (A549^TMPRSS2+^ or A549^CD13+^) cell lines

To generate A549 cells overexpressing CD13 and/or TMPRSS2, cells were transduced with lentiviral vectors harboring sequences for CD13 and/or TMPRSS2. The medium during transduction was supplemented with 8 µg/ml of polybrene (Sigma-Aldrich, Poland) to enhance the process. The medium was refreshed after 16 h, and 72 h later the cells were transferred to a fresh medium that contained blasticidin S (10 µg/ml) and/or G418 (500 µg/ml).

Following this, we implemented clonal selection by plating statistically one cell per well in a 96-well plate. These clones were maintained in DMEM with 20% FBS and selective antibiotics for three weeks to promote proliferation. We targeted clones that achieved over 60% confluence for selection. A relative expression of CD13/aminopeptidase N and TMPRSS2 in each clone was validated through western blotting. Clones with the highest expression of the desired target (CD13 and/or TMPRSS2) were selected for further propagation.

### Virus

We received a clinical isolate of HCoV-229E as a generous contribution from Dr. Lia van der Hoek from Amsterdam University Medical Center, the Netherlands. This isolate was originally procured from an unidentified patient on March 20^th^, 2017 and was propagated oncein HAE, followed by another round of propagation in HuH7 cells. HCoV-229E VR-740 (CCL-171, ATCC) was propagated in MRC-5 cells in DMEM with 2% FBS. Low-passage HCoV-229E clinical isolate (i.e.: passage 2) was used in this study. All HCoV-229E strains were propagated for 96 h at 32°C and 5% CO_2_ before harvesting by a freeze-thaw cycle. Aliquots of viruses were stored at −80°C and the 50% tissue culture infectious dose (TCID_50_) was as described by Reed and Muench (12). The readout was carried out on day 5 p.i.

Before the infection, A549^CD13+^, A549^TMPRSS2+,^ or A549^++^ cells were seeded onto a 96-well plate at 2.5 × 10^4^ cells/well and incubated at 37°C and 5% CO_2_. The culture medium was then removed, and cells were inoculated with 50 µl of 1.4 × 10^4^ TCID_50_/ml of the virus or mock-inoculated. Following an initial 2 h incubation at either 32°C or 37°C, unbound viruses were removed by conducting a minimum of two washes with pre-warmed, serum-free DMEM. Cells were then incubated at either 32°C or 37°C respectively for 5 days unless stated otherwise.

### Viral RNA extraction and Real Time-qPCR

To quantify viral RNA in the supernatant, a 20 µl sample of culture medium was collected, and the sampled volume was replaced with fresh, pre-warmed serum-free DMEM supplemented with P/S. Viral RNA was then isolated from the sample using the MagnifiQTM Viral RNA Kit (A&A Biotechnology, Poland). This isolation procedure was performed on the KingFisher™ Flex Purification System with a 96 PCR head platform (Thermo Fisher Scientific, Poland), following the manufacturer’s instructions.

The quantification of viral RNA was carried out in one-step RT-qPCR using 3 µl of the eluted RNA with the GoTaq^®^ Probe One-step RT-qPCR System (Promega, Poland). This procedure employed *in-house* primers and a probe specifically tailored for the HCoV-229E N sequence: the 229E_NF (sense; 5’-GTTGTGGCCAATGGTGTTAAAG-3’; 600 nM working concentration), 229E_NR (anti-sense; 5’-AGTGTTGCCTGACTCTTTGG-3’; 600 nM working concentration), and 229E_NP (probe; 5’-FAM-ACAATTTGCTGAGCTTGTGCCGTC-TAMRA-3’; 300 nM working concentration).

The reaction was performed on the CFX96™ Touch Real-Time PCR Detection System (Bio-Rad Laboratories, Poland). Cq values were converted into viral RNA copies using a standard curve with a known quantity of viral genome copies. The reaction process consisted of an initial phase of 15 min at 45°C and 2 min at 95°C, followed by 40 cycles of 15 sec at 95°C and 1 min at 60°C.

### Western blot

Protein expression was verified through Western blot analysis of cell lysates. Initially, cells were harvested in a RIPA buffer (comprising 50 mM Tris, 150 mM NaCl, 1% Nonidet P-40, 0.5% sodium deoxycholate, 0.1% SDS; pH=7.5), complemented with 0.5 M EDTA and a protease inhibitors cocktail (cOmplete Tablets, Roche, Poland), and incubated for a minimum of 10 min at 4°C. These cell lysates were mixed with 6×SDS-PAGE sample buffer (0.5 M Tris, 10% SDS, 50 mg/ml DTT; pH=6.8) and denatured at 95°C for 10 min. 30 µg of protein was loaded onto a 4%-12% Bis-Tris gel and separated using SDS-PAGE electrophoresis, in parallel with dual-color PageRuler pre-stained protein size markers (Thermo Fisher Scientific, Poland). The proteins were subsequently electrotransferred onto a polyvinylidene difluoride membrane (PVDF, GE Healthcare, Poland) in a chilled transfer buffer (25 mM Tris, 192 mM glycine, 20% methanol) at 100 V for 70 min. The membranes were blocked with 1× Tris-buffered Saline with Tween-20 (TTBS), supplemented with 5% skim milk (Bioshop, Poland), for 2 h at room temperature (RT). After blocking, membranes were incubated with rabbit anti-human CD13 (monoclonal [EPR4058], 1:2500, Abcam, Poland) and mouse anti-human TMPRSS2 (monoclonal, clone P5H9-A3, cat num: MABF2158, 1:500, MERCK, USA) for a minimum of 2 h at RT, and subsequently washed thrice with TTBS. The membranes were then incubated with either horse radish peroxidase (HRP)-labeled anti-rabbit IgG (1:20000, Agilent Dako, Poland) or HRP-labeled anti-mouse IgG (1:20000, Sigma Aldrich, Poland) antibodies for 1 h at RT. All antibodies were diluted in TTBS with 2.5% skim milk.

Immediately before signal development, membranes were treated with Pierce™ ECL Western blotting substrate under the manufacturer’s protocol (Thermo Fisher Scientific, Poland). Signals were then developed on the membranes using the ChemiDoc Imaging System (Biorad, Poland). Lysates of in-house HAE culture were included as control and equal amount of protein was loaded and processed for analyses.

### Immunostaining

Confluent cells were seeded on coverslips 48 h before infection. Subsequently, cells were infected with 50 µl of HCoV-229E clinical isolate at 1.4 × 10^5^ TCID_50_/ml and incubated for 48 h. Cell monolayers were then washed with 1× PBS and fixed in 4% paraformaldehyde (Chempur, Poland) for 15 mins at RT. Fixed cells were washed thrice in 1×PBS, followed by permeabilization in 0.5% TRITON-X (Sigma Aldrich, Poland) for 13 min at RT. Subsequently, cells were rinsed again thrice with 1×PBS before an overnight blocking stage at 4°C in 1×SEA BLOCK blocking buffer (Thermo Fisher Scientific, Poland).

Cells were then incubated overnight at 4°C with rabbit anti-HCoV-229E nucleocapsid (1:400, Sino Biological) diluted in 1×SEA BLOCK, followed by triple washing with 1× PBS. Next, cells were incubated for 2 h at RT with AlexaFluor (AF) 488-labeled donkey anti-rabbit antibody (1:500, Invitrogen, Poland) and AF647-labeled phalloidin (1:400, Invitrogen, Poland), both diluted in 1×SEA BLOCK.

Afterward, cells were washed thrice with 1×PBS and incubated for 15 min at RT with 0.1 µg/ml of 4’,6-diamidino-2-phenylindole (DAPI, Sigma Aldrich, Poland). After a final three washes with 1×PBS, cells were treated with ProLong™ Diamond Antifade Mountant (Invitrogen, Poland) and mounted for microscopic analyses. Images were acquired under a Zeiss LSM 710 confocal microscope (Carl Zeiss Microscopy GmbH; release version 8.1) using ZEN 2012 SP1 software (Carl Zeiss Microscopy GmbH; black edition, version 8.1.0.484). Images were processed using the ImageJ FIJI version (National Institutes of Health, Bethesda, MD, USA) and are presented as the maximal projections unless stated otherwise.

### Inhibition of entry

Confluent cells were seeded in 96-well plates, and subsequently preincubated at 32°C for 1 h in 50 µl of serum-free DMEM. The medium contained one of the following inhibitors at twice its working concentration: camostat mesylate (500 µM, Sigma Aldrich), E64d (100 µM, Sigma Aldrich), ammonium chloride (10 mM, Sigma Aldrich), or bafilomycin A1 (10 nM, Sigma Aldrich).

Following preincubation, 50 µl of the virus at 2.8 × 10^4^ TCID_50_/ml was added to cells, resulting in a working inoculum of 1.4× 10^4^ TCID_50_/ml and 1× working concentration of respective inhibitors. Cells were subsequently incubated for 2 h at 32°C, after which the inoculum was removed and cells were washed twice with pre-warmed, serum-free DMEM. After that, cells were incubated at 32°C until 48 h p.i in a serum-free medium with the indicated concentration of inhibitors. 20 µl of culture medium was collected to quantify viral RNA by RT-qPCR. To simultaneously assess the cytotoxicity of inhibitors, 96-well plates containing confluent cells were similarly preincubated for 1 h at 32°C in 50 µl of inhibitors at twice the intended concentration. Subsequently, 50 µl of the serum-free medium was added, and the cells were incubated for 2 h. After removing the medium and washing the cells twice with pre-warmed, serum-free DMEM, the cells were maintained for 48 h at 32°C in a serum-free medium containing the specified concentration of inhibitors. Post-incubation, the medium was discarded, and the cytotoxicity of inhibitors was evaluated using the Cell Proliferation Kit (XTT-based, Biological Industries) as per the manufacturer’s protocol. Untreated and DMSO-treated cells were included as controls.

### Statistical analyses

Statistical analyses were performed on GraphPad Prism 6 software. Gaussian distribution test was carried out and data are presented as the median of a triplicate of each sample ± IQR due to the non-normality of data distribution. Comparison of differences in viral RNA copy number and log_10_ transformed viral titer was carried out using the Kruskal-Wallis test followed by a Dunn’s multiple-comparison test. P-values of <0.05 were considered significant.

## Results

### Development of CD13 and TMPRSS2 Overexpressing A549 Cell Lines

To create a robust and stable human alveolar cell line suitable for HCoV-229E studies, A549 cells were co-transduced with lentiviruses carrying the human CD13 and TMPRSS2 genes. These transduced cells underwent selection with G418 and/or blasticidin S. Following clonal selection, the expressions of CD13 and TMPRSS2 in each monoclonal line were assessed, with clones demonstrating the highest expression of CD13 and/or TMPRSS2 selected for further examination. Western blot analyses using human CD13-specific antibodies revealed a single band at approximately 160 kDa only from A549^CD13+^ and A549^++^ cell lysates (**Figure 1A**). Whereas, analyses using human TMPRSS2-specific antibodies revealed two bands at approximately 55 kDa and 60 kDa from A549^TMPRSS2+^ and A549^++^ cell lysates (**Figure 1B**). Furthermore, an additional band at 25 kDa was also detected from the A549^++^ cell lysate (**Figure 1B**). In contrast to the engineered A549 cell lines, only a singular band at approximately 40 kDa was detected from HAE lysates from the application of a human TMPRSS2-specific antibody (**Figure 1B**).

**Figure 1.**
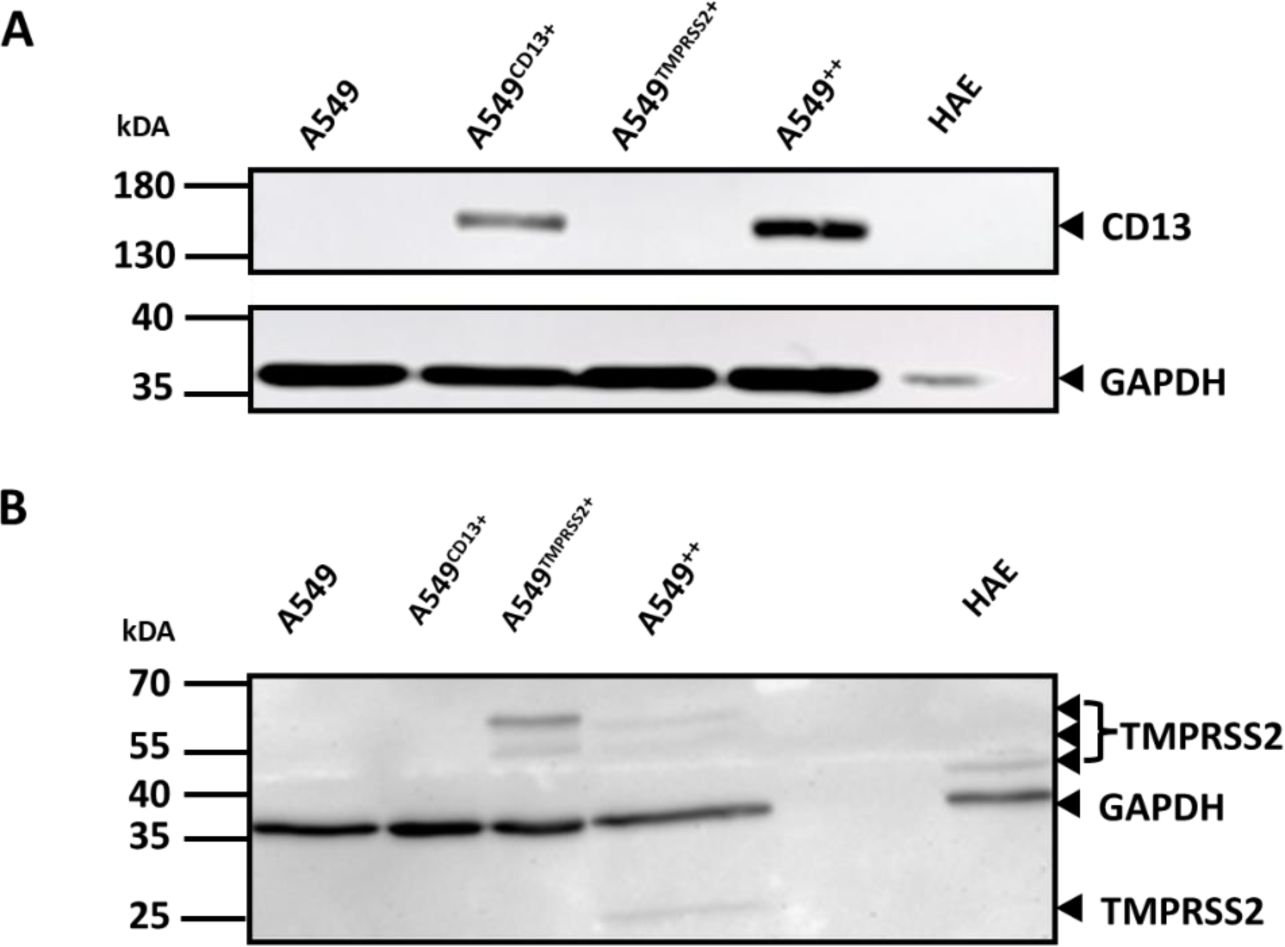
Expression of CD13 and TMPRSS2 in engineered A549 cells. 30 µg of protein from cell lysates was loaded onto a 4%-12% Bis-Tris gel and separated by SDS-PAGE before expressions of (A) CD13 and (B) TMPRSS2 were analyzed by Western Blot. GAPDH detection is included as the loading control. WT A549 cells are included as a negative control, whereas HAE lysates are included as a positive control for TMPRSS2

### Replication kinetics of HCoV-229E in A549^++^ cells

To determine the permissiveness of A549^++^ cells to HCoV-229E, we inoculated A549^CD13+^, A549^TMPRSS2+^, WT A549, or A549^++^ cells with either a clinical isolate of HCoV-229E or HCoV-229E VR-740 and evaluated the viral RNA replication kinetics in the supernatant using RT-qPCR, up to 120 h p.i at 32°C or 37°C. We compared the replication kinetics for A549^++^ cells with those of A549^CD13+^, A549^TMPRSS2+^, and WT A549 cells. We observed a sharp increase in HCoV-229E *N* RNA copy number in culture supernatants from A549^++^ cells by 48 h p.i at 32°C, compared to single-transduced and WT A549 cells **(Figures 2A-F, left panel)**. However, at 37°C, the increase in virus yield was relatively marginal **(Figures 2A-F, right panel)**. Simultaneously, immunostaining of cells infected with the HCoV-229E clinical isolate at 48 h p.i. demonstrated extensive syncytia formation and disruptions in the cell monolayer only in A549^++^ cells at 32°C (**Figure 2G**).

**Fig 2.**
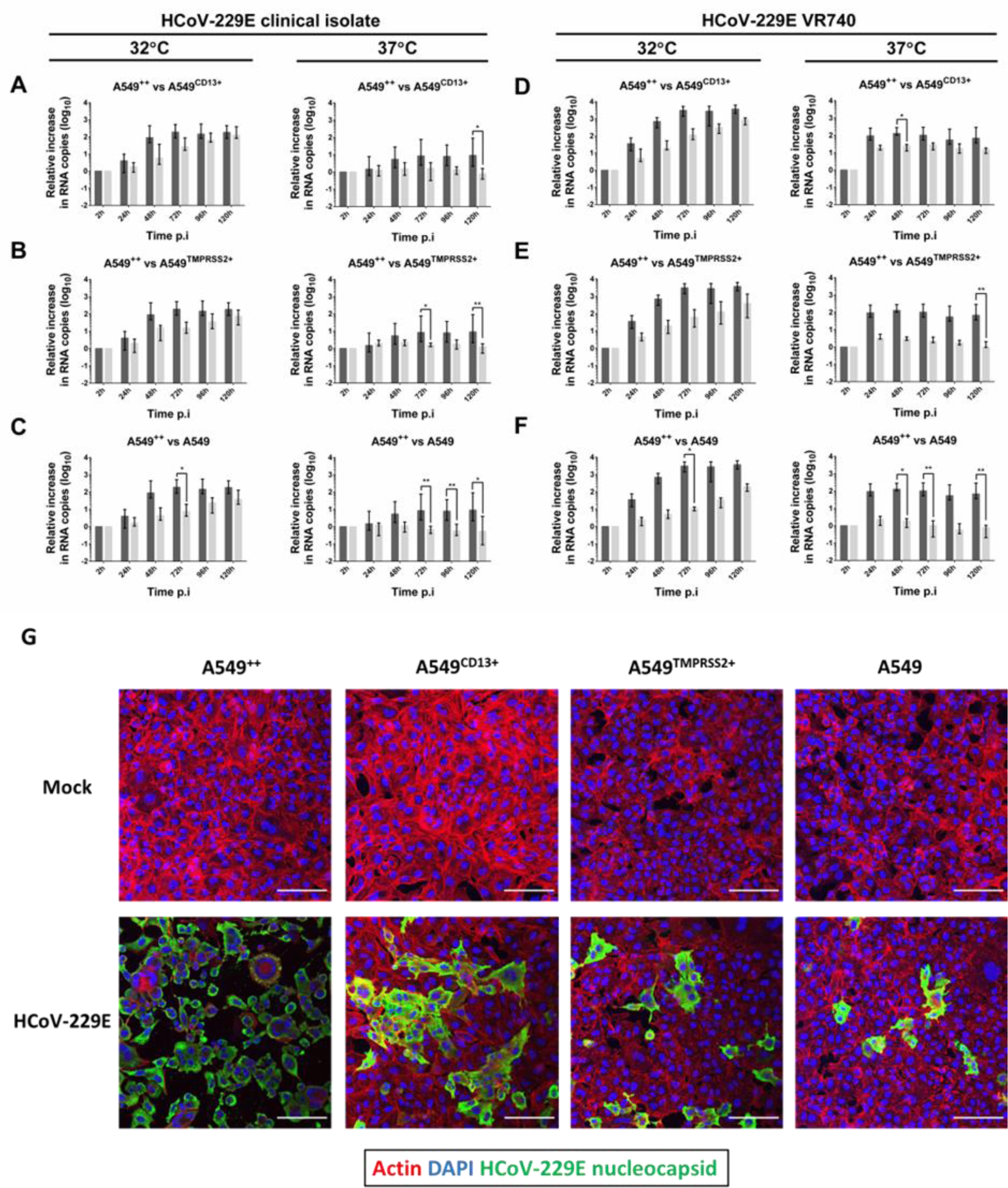
Replication of HCoV-229E on A549^++^ cells. Cells were infected with 1.4 × 10^4^ TCID_50_/ml of HCoV-229E clinical isolates or HCoV-229E VR-740 at 32°C or 37°C. Graphs represent the relative change in the amount of viral RNA in the supernatant over time, whereby relative change in *N* RNA copy number of HCoV-229E clinical isolate in A549^++^ cells (black bars) was compared with that of **(A)** A549^CD13+^, **(B)** A549^TMPRSS2+^, and **(C)** normal A549 cells (all grey bars) for up to 120h p.i. at 32°C and 37°C. Similarly, the relative change in N RNA copy number of HCoV-229E VR-740 in A549^++^ cells (black bars) was also compared with that of **(D)** A549^CD13+^, **(E)** A549^TMPRSS2+^, and **(F)** normal A549 cells (all grey bars). **(G)** Infected cells were fixed at 48h p.i and stained for HCoV-229E nucleocapsid (green) and actin (red). Nuclei were stained with DAPI in blue. The values represent the log_10_ increase in median HCoV-229E *N* RNA copy number relative to 2h p.i. Error bars represent IQR. Data were acquired from two independent experiments with N = 6 for each sample. No asterisk = no statistical significance. *p < 0.05; **p < 0.01, ***; p < 0.001; ****p < 0.0001.

### Infectious HCoV-229E Progeny Production in A549^++^ Cells

To verify whether infected A549^++^ cells produce infectious HCoV-229E progeny, the collected cell culture supernatants were titrated on A549^++^ cells. At 32°C, a significant rise in viral titer was detected in A549^++^ cells infected with the HCoV-229E clinical isolate by 72 h p.i. (**Figure 3A-C, left panel**), whereas the viral titer of HCoV-229E VR-740 plateaued by 48 h p.i (**Figure 3D-F, left panel**). At 37°C, only a mild increase in viral titer was observed in A549^++^ cells infected with the HCoV-229E clinical isolate, which plateaued off by 48 h p.i. (**Figure 3A-C, right panel**). Similarly, the viral titer for the HCoV-229E VR-740 strain plateaued by 72 h p.i. (**Figure 3D-F, right panel**).

**Figure 3.**
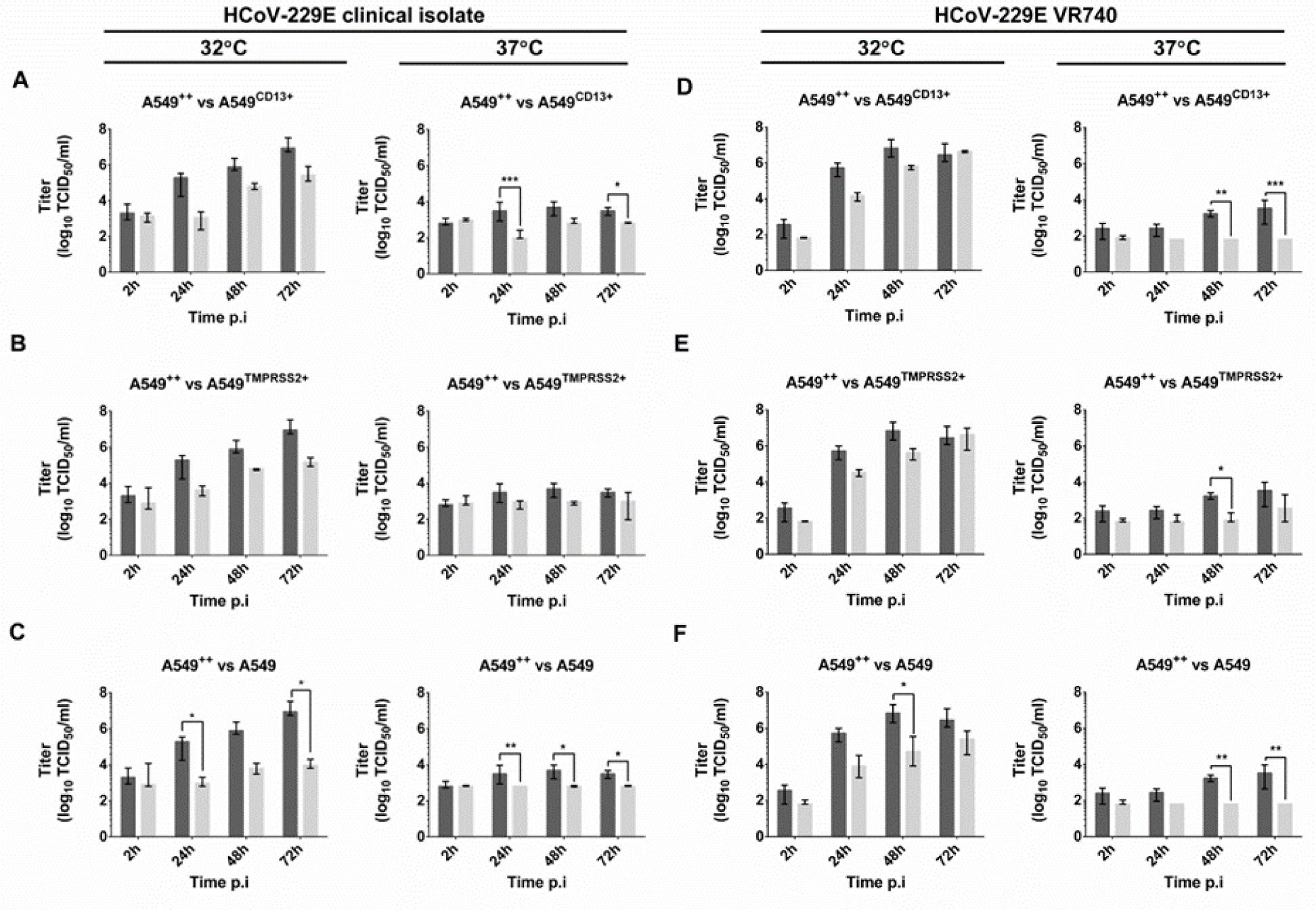
Infectious virus titer kinetics of HCoV-229E in A549^++^ cells. Cells were infected with 1.4 × 10^4^ TCID_50_/ml of HCoV-229E clinical isolate or HCoV-229E VR-740 at 32°C or 37°C. Supernatants from infected A549^++^ (black bar), A549^CD13+^, A549^TMPRSS2+^, and A549 cells (grey bars) up to 72 h p.i were serially diluted and inoculated onto fresh A549^++^ cells. Inoculated cells were incubated for 5 days at 32°C before viral titer readout. HCoV-229E clinical isolate titer kinetics in A549^++^ cells was compared with that of (A) A549^CD13+^, (B) A549^TMPRSS2+^, and (C) normal A549 cells for up to 72 h p.i. at 32°C and 37°C. Similarly, HCoV-229E VR-740 titer kinetics in A549^++^ cells was compared with that of (D) A549^CD13+^, (E) A549^TMPRSS2+^, and (F) normal A549 cells. The values represent the median viral titer, with error bars representing IQR. Data are acquired from two independent experiments with N = 6 per sample. No asterisk = no statistical significance. *p < 0.05; **p < 0.01; ***p < 0.001; ****p < 0.0001.

### Mapping the entry routes of HCoV-229E clinical isolate

WT human coronaviruses have been shown to prefer the TMPRSS2-mediated membrane fusion entry pathway compared to the endocytic pathway, while the cell-culture-adapted variants preferred the endocytic pathway (3, 4, 7). Using the A549^++^ cells, we tested known inhibitor of surface serine proteases (e.g., camostat mesylate) or an inhibitor of endosomal proteases (E64d). The latter category included those that directly impede cysteine proteases like cathepsins B or L or those that inhibit pH-dependent endocytosis, such as the lysosomal acidification inhibitor, ammonium chloride, and the phagolysosome inhibitor, bafilomycin A1. We monitored the cytotoxicity of the compounds (**Figure 4A**) and virus replication kinetics of clinical isolate with RT-qPCR for 48 h p.i. We observed the inhibition of replication in camostat mesylate-treated A549^++^ cells, however, the effect was stronger when camostat and E64d were added together (**Figure 4B**). In contrast, a complete inhibition of viral replication was recorded in E64d-treated A549^CD13+^ cells (**Figure 4C**). Co-application of camostat mesylate and E64d abrogated viral replication in both A549^++^ and A549^CD13+^ cells (**Figure 4B-C**). Taken together, the data indicate that, although HCoV-229E prefers TMPRSS2-mediated entry, it can also readily use a cathepsin-mediated pathway in the absence of TMPRSS2.

**Figure 4.**
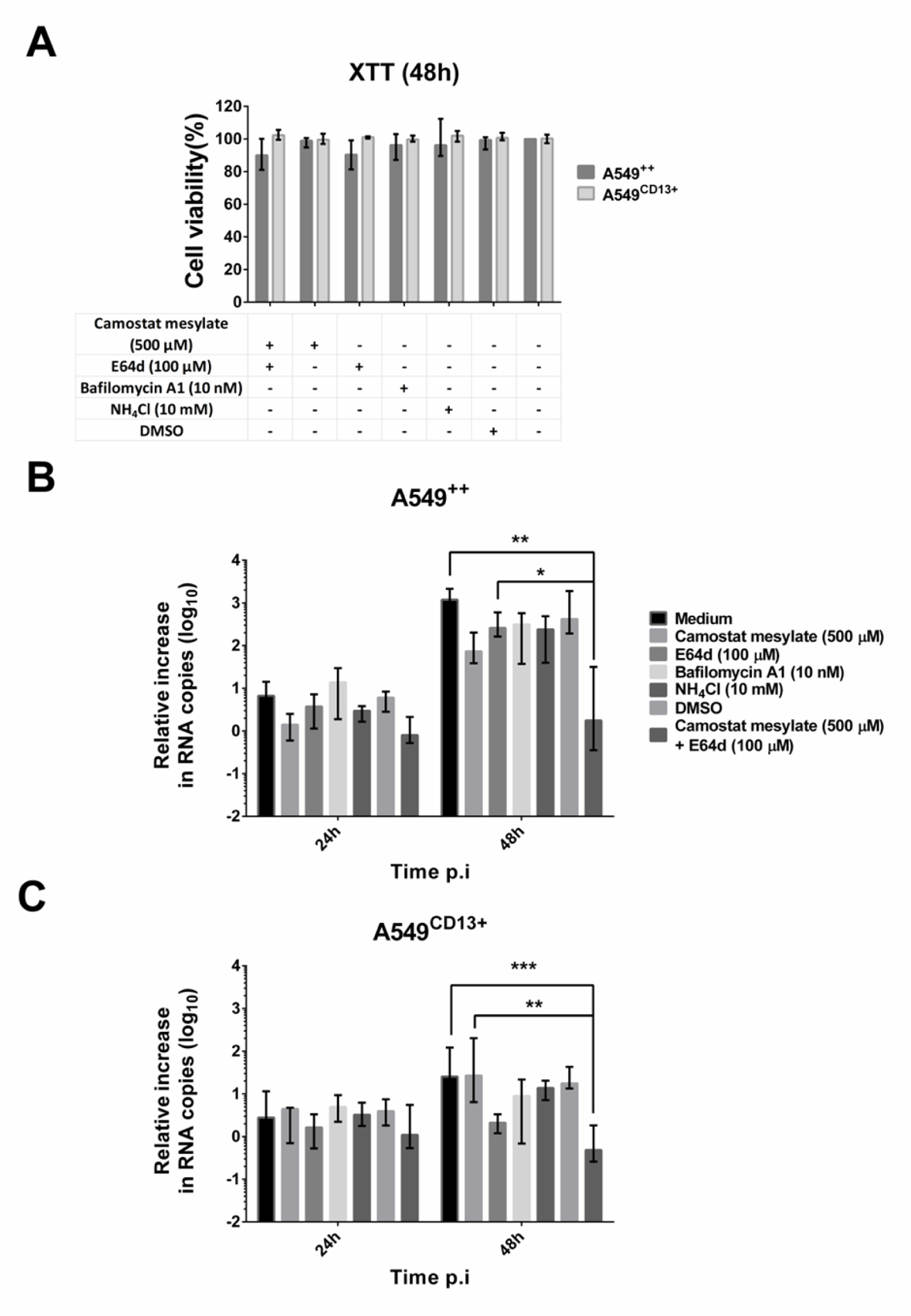
Mapping HCoV-229E clinical isolate entry route. A549^++^ cells were pre-incubated with one of camostat mesylate, E64d, bafilomycin A1, ammonium chloride (NH_4_Cl), or a combination of camostat mesylate and E64d, before infection with HCoV-229E clinical isolate at 32°C for 2h. The supernatant was extracted at 2h-, 24h-, and 48h p.i to measure viral RNA yield by qPCR, and cells were maintained in a medium with respective inhibitors throughout. (A) represents the relative viability of A549^++^ and A549^CD13+^ cells from the application of inhibitors. The values represent the median percentage of viable cells with error bars representing IQR. (B) and (C) represent viral RNA yield in the supernatants of treated A549^++^ cells and A549^CD13+^ cells respectively. The values represent the median relative increase in RNA copy number normalized to 2 h p.i., with error bars representing IQR. Data are acquired from two independent experiments with N = 6 per sample. No asterisk = no statistical significance. *p < 0.05; **p < 0.01; ***p < 0.001; ****p < 0.0001.

## Discussion

Stem cell-derived models of human airway epithelia are widely employed to study coronavirus infections *ex vivo* (13–15). While these models closely mimic the composition and physiology of host tissues, their development is both time-intensive and costly, requiring substantial cell culture expertise (5, 6). These limitations make them less viable for high-throughput applications. Meanwhile, the majority of cell lines are non-permissive to human coronavirus clinical isolates. To address this issue, we developed a robust lung-derived A549 cell line with strong expression of CD13 and TMPRSS2 that crucially, facilitates HCoV-229E clinical isolate replication. While the existing cell lines expressing TMPRSS2, such as HeLa^TMPRSS2+^ (3, 16), have proven effective for studying WT HCoV-229E infections as an alternative to HAE, they are less suitable for pathological studies due to their non-lung tissue origin and defective interferon response (17, 18). We believe that our lung-derived A549 model offers a more physiologically relevant *in vitro* platform for studying human coronavirus infections.

Our findings affirm the significance of both CD13 and TMPRSS2 for HCoV-229E clinical isolate infection. Consistent with previous research (4), we observed that syncytia formation in HCoV-229E-infected cells is enhanced when both CD13 and TMPRSS2 are present in the host cell. It has been demonstrated that high CD13 density enhances porcine epidemic diarrhea virus (PEDV) infection (19). Similarly, TMPRSS2 has been found to amplify viral replication and syncytia formation in HCoV-NL63, MERS-CoV, and SARS-CoV-2 infections (20–22). Altogether, our results highlight that an effective infection by HCoV-229E clinical isolate necessitates both viral spike priming by TMPRSS2 and optimal host cell expression of CD13.

In contrast to previous observations, our data indicate that in the absence of TMPRSS2, the HCoV-229E clinical isolate can still gain cell entry using a cathepsin-mediated phagolysosomal pathway. A previous study by Shirato et al suggested that HCoV-229E clinical isolates need multiple passages to develop the capability for endosomal entry in standard HeLa cells (7). In our study, we employed a low-passage HCoV-229E clinical isolate, and it remains uncertain whether our observations merely reflect differences in the spike sequence across varying HCoV-229E clinical strains. However, recent research into the cellular entry mechanisms of SARS-CoV-2 variants suggests that the Omicron variant has a different preference for activating proteases, as compared to previous variants (23, 24). These findings could signify an evolutionary adaptation of coronaviruses to enhance cellular entry efficiency.

During the model development, we observed that TMPRSS2 is produced in two forms (55 kDa and 60 kDa), in contrast to the native HAE model. However, the TMPRSS2 species correspond in size to the putative full-length zymogen and an *N-*glycosylated zymogen (25, 26). The additional protein species (25 kDa) observed only in A549^++^ cells, on the other hand, is consistent with the approximate size of the TMPRSS2 S1 peptidase domain (27). This suggests that overexpression of CD13 in A549^++^ cells may contribute to the activation of TMPRSS2 and the shedding of the S1 peptidase domain. In HAE however, we only observed one TMPRSS2 species at 40 kDa. The presence of the 40 kDa TMPRSS2 species had been noted previously in both human semen and Lymph Node Carcinoma of the Prostate (LNCaP) cell line under boiling and non-reducing conditions, indicating that TMPRSS2 forms distinct molecular complexes from tissue to tissue (25).

In conclusion, we have developed a robust and physiologically relevant cell line model that is permissive to HCoV-229E clinical isolate replication. Our data on the viral replication kinetics, cellular pathology, infectious progeny production, and entry inhibition assay highlight the potential of A549^++^ cells for further studies on HCoV-229E clinical isolate infection *in vitro.* Furthermore, our data further support what has been known about HCoV-229E infectivity and provide further insight into the potential of lentiviral transduction in developing permissive cell models for viral infection studies.

## CRediT authorship contribution statement

### Laurensius Kevin LIE

Conceptualization, Methodology, Validation, Investigation, Formal analysis, Visualization, Writing - original draft.

### Aleksandra SYNOWIEC

Conceptualization, Methodology, Validation, Investigation, Formal analysis, Visualization, Writing - review & editing.

### Jedrzej MAZUR

Investigation, Methodology.

### Krzysztof PYRĆ

Conceptualization, Methodology, Validation, Writing - review & editing, Resources, Supervision, Funding acquisition.

